# Atmospheric transport of particulate matter and particulate-bound agrochemicals from beef cattle feedlots: human health implications for downwind agricultural communities

**DOI:** 10.1101/2023.03.22.533817

**Authors:** Amanda D. Emert, Frank B. Green, Kerry Griffis-Kyle, Philip N. Smith

## Abstract

**Background:** Beef cattle feedlot-derived particulate matter (PM) is a complex mixture of dust, animal waste, agrochemicals, and bioaerosols. No empirical data currently exists quantifying human exposure of PM-bound agrochemicals downwind of feedlots.

**Objectives:** There were three objectives of the current study: 1) to determine spatial extent and magnitude of PM transport downwind of large beef cattle feedlot facilities, 2) to quantify occurrence of pyrethroid insecticides and anthelmintics in feedlot-derived PM, and 3) to assess cumulative human health risk of agrochemicals in agriculture-adjacent communities downwind of feedlots.

**Methods:** Authors investigated downwind transport (<1 to >12 km) of total suspended particulates (TSP) from three feedlots in the Southern Great Plains (SGP) of North America. PM collected on TSP filters was analyzed via UHPLC-MS/MS for six pyrethroids (bifenthrin, λ-cyhalothrin, cypermethrin, esfenvalerate, fenvalerate, and permethrin) and five macrocyclic lactones (MLs; abamectin, doramectin, eprinomectin, ivermectin, and moxidectin). An empirical distance decay model was used to determine probabilistic PM concentrations in downwind ambient air.

**Results:** Downwind TSP concentrations exhibited rapid decline from 0.01 - ≤1.6 km (Monte Carlo-simulated mean ± SEM; 5,049 ± 96.1 µg/m^3^) and subsequent stabilization >1.6 – 12.4 km (1,791 ± 9.9; µg/m^3^). TSP concentrations did not converge to background levels within the spatial extent of the study (12.4 km). Agrochemicals were detected downwind >LOQ at greater overall frequency (40.6%) than upwind locations (26.8%). Two pyrethroids were detected at the highest overall downwind concentrations (mean ± SEM; fenvalerate = 5.9 ± 0.8, permethrin = 1.1 ± 0.3 ng/m^3^), and screening-level cumulative exposure estimates indicate elevated pyrethroid risk (LOC = 1; RI = 0.173) in children (1-2 yrs) living near commercial agricultural operations in the SGP.

**Discussion:** Results significantly expand the known distribution of feedlot-derived PM and agrochemicals, and consequently highlight exposure pathways unrecognized in residential human health assessments and feedlot risk evaluations.

## 1. Introduction

Confined beef cattle feeding operations, hereafter “feedlots,” have become standard practice in developed countries for rapidly increasing muscle mass and fat cover in beef cattle prior to slaughter (Smith, 2021). Feedlot cattle density can range from hundreds to hundreds of thousands of head contained in open-air, earthen floor pens. In addition to loose soil, pen floor material contains a complex mixture of chemical and biological contaminants, including veterinary pharmaceuticals metabolized and excreted via feces and urine (Aust et al., 2008; Blackwell et al., 2014; Blackwell, Johnson, et al., 2015; Blackwell, Wooten, et al., 2015; Zhao et al., 2010), viral and bacterial pathogens (Alexander et al., 2011; Gaudino et al., 2022; Mosier, 2015; Qian et al., 2018), and pesticides (Modernel et al., 2013; Sadler et al., 2005; Yates et al., 2011). While it is well-documented that these facilities are detrimental to watersheds through off-site runoff of manure (Bicudo & Goyal, 2003; Sahoo et al., 2016), mounting evidence in the last two decades also implicates aerial dispersion of PM as a major transport pathway of agrochemicals beyond feedlot boundaries (Emert et al., 2023; McEachran et al., 2015; Peterson et al., 2017, 2020, 2021, 2022; Sandoz et al., 2018; Wooten et al., 2018). This is particularly problematic in arid and semi-arid regions, where dry pen floor material is aerosolized by cattle movement-driven mechanical disturbance (de Oro et al., 2021), which is then wind-transported downwind. Presently, U.S. EPA does not include particulate-bound emissions as an agrochemical transport pathway in environmental risk evaluation of feedlot facilities (U.S. EPA, 2004) or human health assessments supporting pesticide registration (U.S. EPA, 2012).

Downwind trajectories of suspended particulates are dependent upon atmospheric conditions and individual particle geometry (i.e., size and shape). More broadly, however, coarse PM size fractions (PM_10_; ≤10 - >2.5 μm) and larger debris settle out of suspension rapidly, exhibiting saltatory movement over smaller distances, but fine PM fractions (PM_2.5;_ ≤2.5 – >0.1 μm) are capable of atmospheric transport exceeding hundreds of kilometers (Chen et al., 2014; Jiménez-Vélez et al., 2009; Kallos et al., 2007; Perry et al., 1997). Particulate size fractions are also relevant to human exposure. PM_10_ is capable of deposition in the human respiratory tract, and PM_2.5_ is capable of transport across the alveolar-capillary barrier deep in the lungs (Arias-Pérez et al., 2020). Chronic exposure to inhalable PM is linked to numerous adverse health outcomes including respiratory and cardiovascular diseases, cancers (Group I carcinogen; IARC, 2016), inflammation, adverse birth outcomes and overall lower life expectancies (Mukherjee & Agrawal, 2017). Moreover, susceptibility to PM-associated disease is greater in children and older adults (Sacks et al., 2011). A 2014 meta-analysis of North American and European populations observed 8 and 9% increased incidences of lung cancer per 10 μg/m^3^ increase of outdoor PM_10_ and PM_2.5_ exposure, respectively (Hamra et al., 2014). As such, inhalable PM in ambient air is hazardous to human respiratory health irrespective of its potential as an agrochemical transport vehicle.

Pyrethroid insecticides are the most frequently detected pesticide class in feedlot PM (Peterson et al. 2020) and have largely superseded older chemical classes (carbamates, organophosphates) used for control of external livestock and foliar cropland pests. Pyrethroids are neurotoxic, interacting with voltage-gated sodium ion channels to cause hyperpolarization of neuronal membranes, paralysis, and eventual death in target and non-target invertebrates (Bradberry et al., 2005; Soderlund, 2012; Todd, 2003). Pyrethroid exposure has recently been associated with a higher risk of all-cause and cardiovascular-specific human mortality (Bao et al., 2020), and agricultural workers and their children exposed to pyrethroids have a higher risk of impaired neurofunction (cognitive, motor, and behavioral; Lucero & Muñoz-Quezada, 2021). In humans living or working near pyrethroid pesticide applications in the previous five years, epigenome-wide association has identified differential methylation at CpG sites associated with Alzheimer’s disease, diabetes, birth defects, and several cancers (n = 237; Furlong et al., 2020). Chronic exposure has been linked to primary ovarian insufficiency in Chinese women (Li et al., 2018) and male rat reproductive disfunction (oral; low total mixture dose = 0.0006 mg/kg/day, high total mixture dose = 0.0031 mg/kg/day; Ravula & Yenugu, 2021), including changes in fecundity and testicular genotoxicity (total mixture dose (oral) = 0.016 mg/kg/day; Priyanka et al., 2022). Prenatal exposure to pyrethroid mixtures is associated with adverse effects on human neurodevelopment (thyroid hormone function; Andersen et al., 2022), and irreversible adverse effects on follicular development and ovarian function in female rat offspring (total mixture dose (rat, oral) = 0.2 mg/kg/day; Song et al., 2022).

In addition to pyrethroids, macrocyclic lactones (MLs) including avermectins and milbemycins are the most frequently used veterinary anthelmintics, guarding livestock against both ecto- and endoparasites. Similarly, MLs are neurotoxic, interacting with GABA- and glutamate-gated chloride channels (Lumaret et al., 2012). Mammalian and avian nephrotoxicity, hepatotoxicity, and neurotoxicity are typically expressed in unit magnitudes exceeding mg/kg quantities for oral/dietary exposure (Salman et al., 2022), where 0.006-0.012 mg/kg oral ivermectin and 0.25 mg/kg oral moxidectin represent therapeutic doses for heartworm prevention in dogs (El-Saber Batiha et al., 2020). While human toxicity is typically low, avermectin B1_a,b_ and derivatives are contraindicated for pregnant women due to potential developmental toxicity (avermectin B1; NOAEL = 0.075 mg/kg/day; MADL = 4.4 μg/day; OEHHA, 2011). Repeated sublethal exposure in medically important soil-transmitted helminths may also confer generational ML resistance (Moser et al., 2019; Prichard & Geary, 2019; Tinkler, 2020), an emerging public health concern in veterinary and human medicine (Freeman et al., 2019).

Hence, there were three objectives of the current study: 1) to determine the spatial extent and magnitude of PM transport downwind of large beef cattle feedlot facilities, 2) to quantify occurrence of pyrethroid insecticides and veterinary anthelmintics in feedlot-derived PM, and 3) to assess cumulative human health risk of agrochemicals in agriculture-adjacent communities downwind of feedlots. We first hypothesized that empirical distance decay of total suspended particulates (TSP) measured at three downwind locations could be used to characterize downwind PM transport. Secondly, we hypothesized that particulate-bound agrochemical transport extends beyond the spatial extent reported in previous literature (≤1 km). Through the present objectives, we aim to better inform human and ecological risk assessments of feedlot facilities and feedlot-derived PM.

## 2. Materials and Methods

### 2.1 Site description

The Southern Great Plains (SGP) region of North America encompasses portions of Colorado, Kansas, New Mexico, Oklahoma, and Texas (**Figure 1**). The SGP contains the highest concentration of beef cattle in the U.S. (9.1 million head), centralized in the panhandle of Texas (**Figure 1**; 4.5 million head; 15.6% of US total beef cattle inventory; USDA NASS, 2023). Largely an agroecosystem, the regional burden of agrochemicals used for crop production already poses a risk to terrestrial and aquatic biodiversity (Brausch & Smith, 2009; Emert et al., 2023; Hanberry et al., 2021). Crops grown in the Texas panhandle primarily include cotton (4.9 million ac) and winter wheat (3 million ac), accounting for 70.9% of all cultivated acreage in 2021 (Cropscape Data Layer, USDA NASS, 2021). In addition to mechanical disturbance of manure and exposed topsoil in feedlot pens, windborne soil erosion is considerable in the semi-arid region. The SGP is split along a continental dry line, where warm-dry air from the desert southwest meets warm-moist air from the Gulf of Mexico. Meteorological conditions along the dry line are a primary driver of high winds and tornadic weather systems frequently observed in Texas, Oklahoma, and Kansas in the spring and early-summer season (Hoch & Markowski, 2005). As regional climate changes, prolonged periods of drought are projected to increase (Modala et al., 2017; Shafer et al., 2014), with subsequent increases in natural and anthropogenic PM emissions.

**Figure 1:**
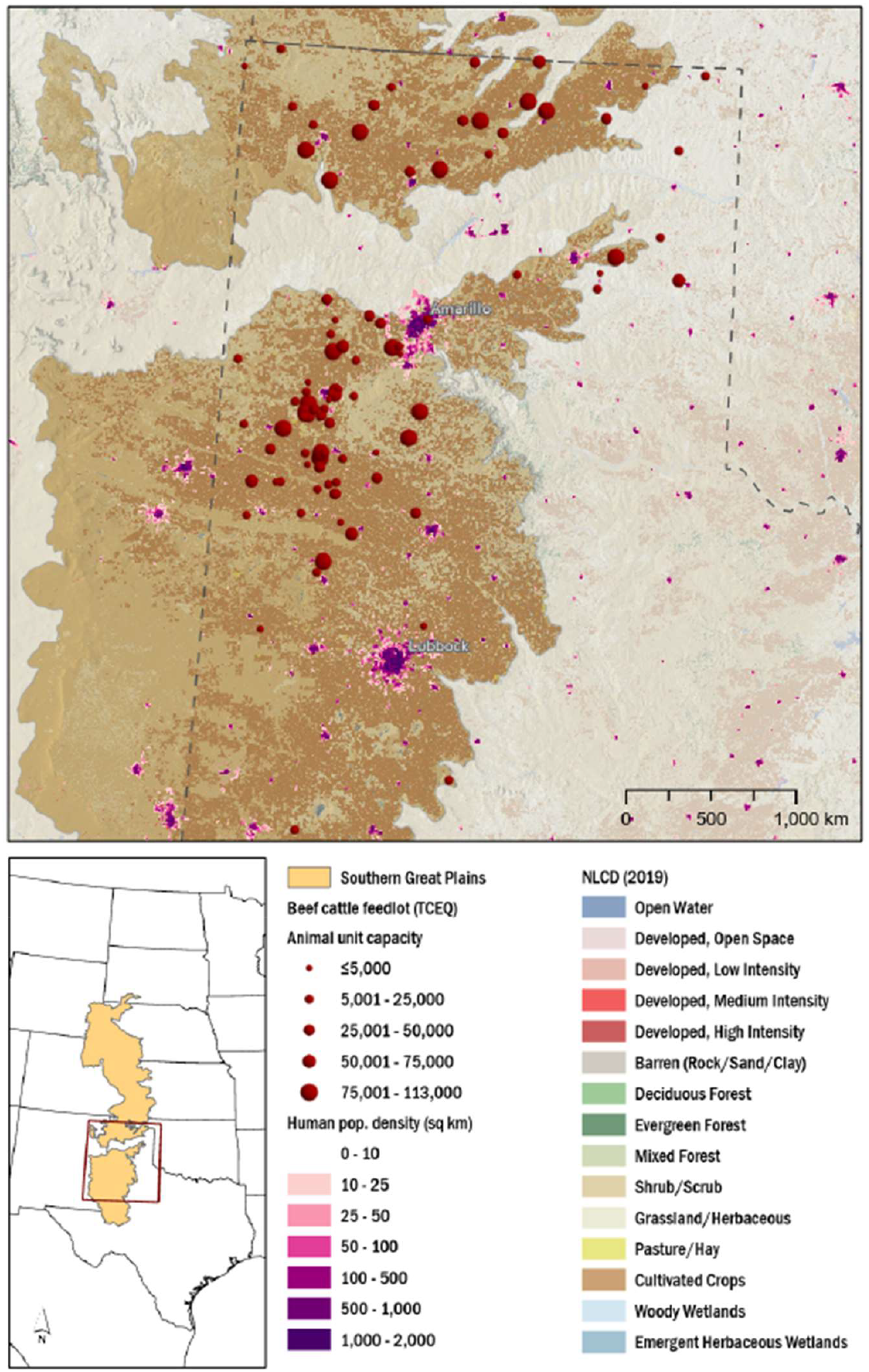
Map of study area in the Southern Great Plains of North America highlighting beef cattle feedlot facilities (TCEQ, 2021), human population density per sq km (WorldPop, 2023), and land cover (MRLC, 2022).

### 2.2 TSP collection

Total suspended particulates were collected upwind and downwind of three feedlots (F1-F3; animal unit capacity = 75,000-82,500; **Supplemental Material SI1**) in the Texas panhandle over six events per feedlot (n = 18) from May - August 2022. To target peak PM emissions, sample collection occurred in the evening 1.5 hrs. before and after dusk on dates where no precipitation occurred at least 72 hrs. prior to collection. One upwind and three downwind locations (D1-D3; >1 – 12.4 km) were selected based on public road access, wind speed and direction. Feedlots were located >20 km from the nearest neighboring facility to minimize upwind PM influence to the extent practicable. Glass fiber filters (10 cm; CF-902; HI-Q Environmental Products, San Diego, CA) were pre-weighed and sealed in air-tight containers and bagged prior to sampling. Four Hi-Q CF-902 digital portable high-volume air samplers (HI-Q Environmental Products, San Diego, CA) were used for TSP collection; two samplers were co-located at each sampling point for 30 min. intervals, with samplers alternated for decontamination (70% ethyl alcohol) between locations. Geographic coordinates were recorded at each sampled location. At the start of each 30 min. interval, weather variables including air temperature and pressure, relative humidity, wind speed and direction were recorded from the nearest West Texas Mesonet station (WTM, 2022). At the end of each sampled interval, filters were placed in containers and bagged on ice until transport to the Texas Tech University Institute of Environmental and Human Health (TIEHH) where they were stored at -20 °C until extraction.

### 2.3 TSP distance decay model validation

A simplified physical model of distance decay was validated and incorporated for the purpose of estimating probabilistic downwind PM concentrations. Modeled TSP distance decay functions were compared using six candidate model distributions (four parameter logistic decay, two and three parameter exponential decay, four and five parameter biexponential decay, and Weibull decay), where the model of best fit was used for training and validation. Training and validation sets consisted of two of three feedlots (F1, F3), stratified by feedlot and downwind sample ID (D1-D3; 50% training: 50% validation). After model validation, predicted bagged mean TSP concentrations (n = 1,000) for all feedlots were calculated, and final model prediction was assessed with the previously unincorporated third feedlot (F2) actual vs. predicted TSP concentrations. Monte Carlo simulations (n = 10,000) were then performed using the validated distance decay function with random noise ≤1 standard deviation from the mean.

### 2.4 Materials

Neat analytical standards (purity; 95-99.5%) were acquired from Sigma-Aldrich (Burlington, MA; abamectin (ABA), cypermethrin (CYP), doramectin (DORA), eprinomectin (EPRIN), ivermectin (IVM), moxidectin (MOX), permethrin (PERM), tris (1-chloro-2-propyl) phosphate (TCPP)) and Chem Service (West Chester, PA; bifenthrin (BIF), λ-cyhalothrin (λ-CYH), esfenvalerate (ESFEN; *S,S*-fenvalerate), fenvalerate (FENV)). Magnesium sulfate and sodium chloride salts, high-performance liquid chromatography (HPLC) grade solvents (acetonitrile, acetone, methylene chloride, methanol, and water) and polytetrafluoroethylene (PTFE) syringe filters (0.45 μm, 30 mm; 0.22 μm, 13 mm) were purchased from Thermo Fisher Scientific (Waltham, MA).

### 2.5 Filter extraction and analysis

Sample extraction methodology was performed as described in Peterson et al. (2021). Briefly, TSP filters were weighed individually, and two filters per sampled location were placed in a 50-mL centrifuge tube to ensure sufficient sample mass for analysis. Filters were spiked with internal standard (TCPP), and 45 mL methylene chloride:acetone (1:1) was added, followed by heated sonication (40 °C) for 1 hr.

Supernatant was then syringe filtered (PTFE; 0.45 μm) into a secondary glass vial, and 40 mL acetonitrile:water (1:1) was added to the centrifuge tube. Tubes were capped and placed on an orbital shaker platform (Model MaxQ 4000, Thermo Scientific, Waltham, MA) for 18 hrs at 350 rpm. Next, tubes were centrifuged (10 min; 3100 rpm) and supernatant was decanted into a second 50-mL centrifuge tube. Magnesium sulfate (4 g) and sodium chloride (1 g) salts were added, and tubes were vortexed for 1 min followed by centrifugation for 10 min at 3100 rpm. The final supernatant was then combined with the initial supernatant in the glass vial, evaporated under a stream of nitrogen at 35 °C, and reconstituted in 1 mL acetonitrile. Extract was vortexed for 10 s and syringe filtered (PTFE; 0.22 μm) into amber chromatography vials for subsequent analysis via triple quadrupole liquid chromatography tandem mass spectrometry with electrospray ionization (Thermo TSQ Quantum Access Max, Thermo Scientific, Waltham, MA), as described in Peterson et al. (2020).

### 2.6 Quality assurance/quality control

Trip blanks and field blanks were collected once per day; trip blank filters remained sealed in coolers for the duration of each event, and field blanks were exposed to ambient air for 30 s. Matrix spikes and method blanks (approximate ratio; 10:1) were co-extracted, and additional clean filters were extracted for matrix-matched calibration standards. All analyte calibration curves were above minimum acceptable limits of linearity (R^2^ >0.995; RSD ≤20%), and a signal-to-noise (S/N) ratio of 3:1 was used for quantitation of sample peaks. Limit of detection (LOD) and limit of quantitation (LOQ) were defined as 3.3σ/S and 10σ/S, respectively (σ = standard deviation of detector response; S = slope of calibration curve; ICH, 2021). No analytes were detected >LOQ in any trip blanks or field blanks. All samples were analyzed <5 months of collection.

### 2.8 Statistical Analysis

Prior to multivariate analysis and probabilistic assessment, downwind TSP and agrochemical concentrations were normalized by upwind concentrations (denoted, hereafter, as ΔU), and transformed. If approximate normality of quantile residuals could not be achieved after transformation, non-parametric Friedman’s with Bonferroni correction for multiple comparisons was performed with categorical distance classes (upwind, D1-D3). Analytes were excluded if detections >LOQ were <30%. Where analytical detections were >30%, non-detects were imputed as ½ LOD. Multicollinearity was assessed with robust maximum likelihood (RML) estimation, and multivariate outliers were excluded if they exceeded the upper control limit (UCL) of Mahalanobis’ distance or were identified as residual quantile outliers (**Supplemental Material SI2)**. All predictor and response variables were then centered and scaled. Two levels of multicollinearity were addressed with partial least squares analysis (PLS), such that potential associations between diurnal weather gradients and intraseasonal weather gradients could be determined independently. K-fold cross-validation (k = 7) was employed to determine optimal number of model factors in each case. Retained factor scores from PLS were then incorporated with explanatory variables in canonical correlation analysis (CCA) for repeated measures. Empirical model selection of TSP distance decay was based on five selection parameters (corrected Akaike’s Information Criterion (AICc), sum of square error (SSE), root mean square error (RMSE), and correlation coefficient (R^2^). The best fit model function was used to perform Monte Carlo simulations (n = 10,000), and simulated values were backtransformed for interpretation. Daily weather (stations; n = 3) and wind rose observations (stations; n = 4) from May 1 - August 31, 2022 were obtained from Weather Underground and NOAA Automated Surface Observing Systems (ASOS), respectively, where weather summaries, wind speed and directional probabilities from each station were collated. Parametric statistical analysis, model validation, and uncertainty analyses were performed in JMP Pro v16.0.0 (SAS Institute Inc.; Cary, NC). Non-parametric Friedman’s tests were performed in IBM SPSS Statistical Software for Windows (Version 28.0. Armonk, NY; IBM Corp.)

### 2.7 Cumulative human health assessment

Cumulative short-term (non-carcinogenic) adult (≥16 yrs) and child (1-2 yrs) health assessments and aggregate adult lifetime (carcinogenic) health assessments of relevant agrochemicals were performed, where applicable, incorporating the most recent U.S. EPA screening-level cumulative (U.S. EPA, 2012) aggregate human health assessments (U.S. EPA, 2016, 2017b, 2019a, 2020b, 2020a, 2020c), and SOPs (U.S. EPA, 2005, 2012) as primary sources of methodology and exposure estimates. Feedlot-derived exposure scenarios were limited to parameters of the current study design, assuming 99.9^th^ percentile of seasonal inhalation exposure from a single area source. Cumulative exposure considered all relevant dietary (99.9^th^ percentile), oral, dermal, and inhalation exposure in adults and children (1-2 yrs) in SGP agricultural communities situated downwind of feedlot operations.

Daily inhalation dose (ADD; mg/kg/day; **Eq. 1**) or lifetime-adjusted average daily inhalation dose (LADD; mg/kg/day; **Eq. 2**) were calculated as:

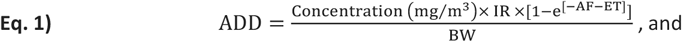

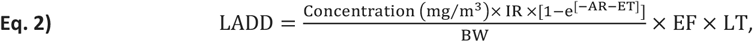

where IR = inhalation rate (m^3^/hr), AF = absorption factor, BW = body weight (kg), ET = exposure time, and LT = lifetime (years). Additionally, a cancer potency factor (Q ^*^) was applied to LADDs (**Eq. 3**) to estimate incremental lifetime cancer risk (ILCR):

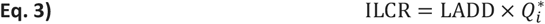

Single compound margin of exposure (MOE; **Eq. 4**) was then calculated individually as:

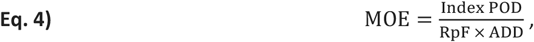

where index POD = point-of-departure of index compound and RpF = relative potency factor. Cumulative risk indices (RI) were then calculated from U.S. EPA dietary, oral, dermal, and inhalation ADD (**Eq. 5**) as:

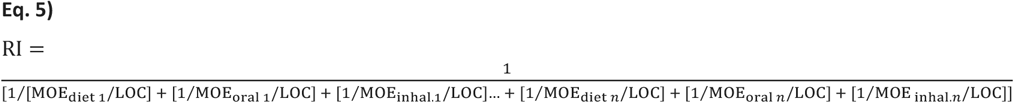

Levels of concern (LOCs) are based on lifestage-applicable uncertainty (UF_A_, UF_H_) and safety factors (FSQA SF; U.S. EPA, 2019), where cumulative agrochemical RI’s >1 are unlikely to pose a human health risk to relevant subpopulations. Generic U.S. EPA recommended values (U.S. EPA, 2017) were applied for body weight (kg), inhalation rate (m^3^/hr), averaging time (AT; adult, non-carcinogen), and lifetime (LT; adult, carcinogen). Exposure parameter values for adults and children (1-2 yrs) are listed in **Table 1**.

**Table 1:** Residential human health exposure parameters

| Parameters | Adult | Child, 1-2 yrs |
| --- | --- | --- |
| Body weight (kg)<sup>a</sup> | 80 | 11 |
| Inhalation rate (m<sup>3</sup>/hr)<sup>a</sup> | 0.64 | 0.33 |
| Exposure time (ET; hrs/day) | 8 | 8 |
| Exposure frequency (EF; days/year; seasonal) | 123 - *x wet days* | NA |
| Exposure duration (ED; life expectancy; years)<sup>b</sup> | 78 | NA |
| Averaging time (years; non-cancer)<sup>b</sup> | 78 | NA |
| Lifetime (LT; years; cancer)<sup>b</sup> | 78 | NA |

<sup>a</sup>U.S. EPA generic lifestage values (U.S. EPA, 2012) <sup>b</sup>U.S. EPA post-application residential values (U.S. EPA, 2011) NA = not applicable

## 3. Results and Discussion

Non-normalized ambient air TSP concentrations (µg/m^3^) are presented in **Figure 2**. Summary statistics of non-normalized TSP concentrations and feedlot proximity are available in **Supplemental Material SI3**.

**Figure 2:**
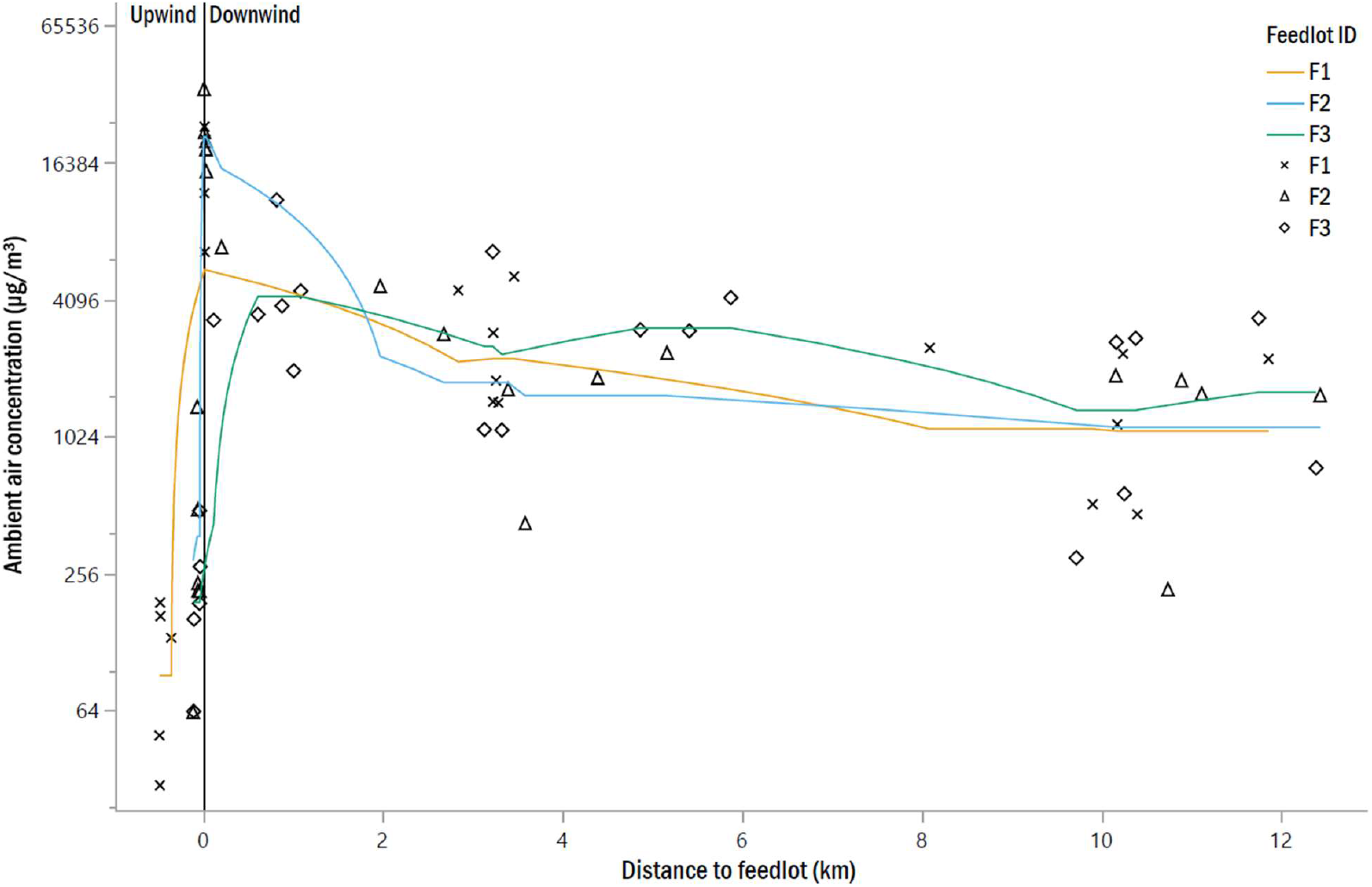
Moving 5-point average of total suspended particulate (TSP) ambient air concentrations (µg/m^3^) upwind and downwind of three feedlots (F1-F3) in the Southern Great Plains, Texas, USA.

### 3.1 PM distance decay and Monte Carlo simulation

Monte Carlo-simulated distance decay serves to highlight the extent to which dust and organic debris remain airborne downwind of feedlots at a height of 2 m. Following model comparison (**Supplemental Material SI4**), a Weibull-distributed ΔU TSP distance decay function (**Eq. 6**) was selected as the most appropriate model,

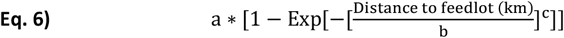

where a = the asymptote, b = the inflection point, and c = the growth rate of the decay curve. Final model fit of actual vs. predicted TSP concentrations and Weibull distance decay curve are presented in **Supplemental Material SI5a-b**. After the model was validated and tested (**Supplemental Material SI6a-b**), the Weibull-distributed function was used to simulate Monte Carlo probabilistic point-density distribution presented in **Figure 3**. Initial and simulated box-cox (λ = -0.109) transformed ΔU TSP and back transformed simulated ΔU TSP parameter estimates are presented in **Table 2**. The current distance decay model elucidates a rapid decline in TSP concentrations from 0 – 1.6 km (simulated mean ± SEM from 0 – 1.6 km = 5,049 ± 96.1; 95% CI: 4,861 – 5,238 µg/m^3^) and subsequent trajectory stabilization >1.6 - 12.4 km (1,791 ± 9.9; 95% CI: 1,773 – 1,811 µg/m^3^).

**Figure 3:**
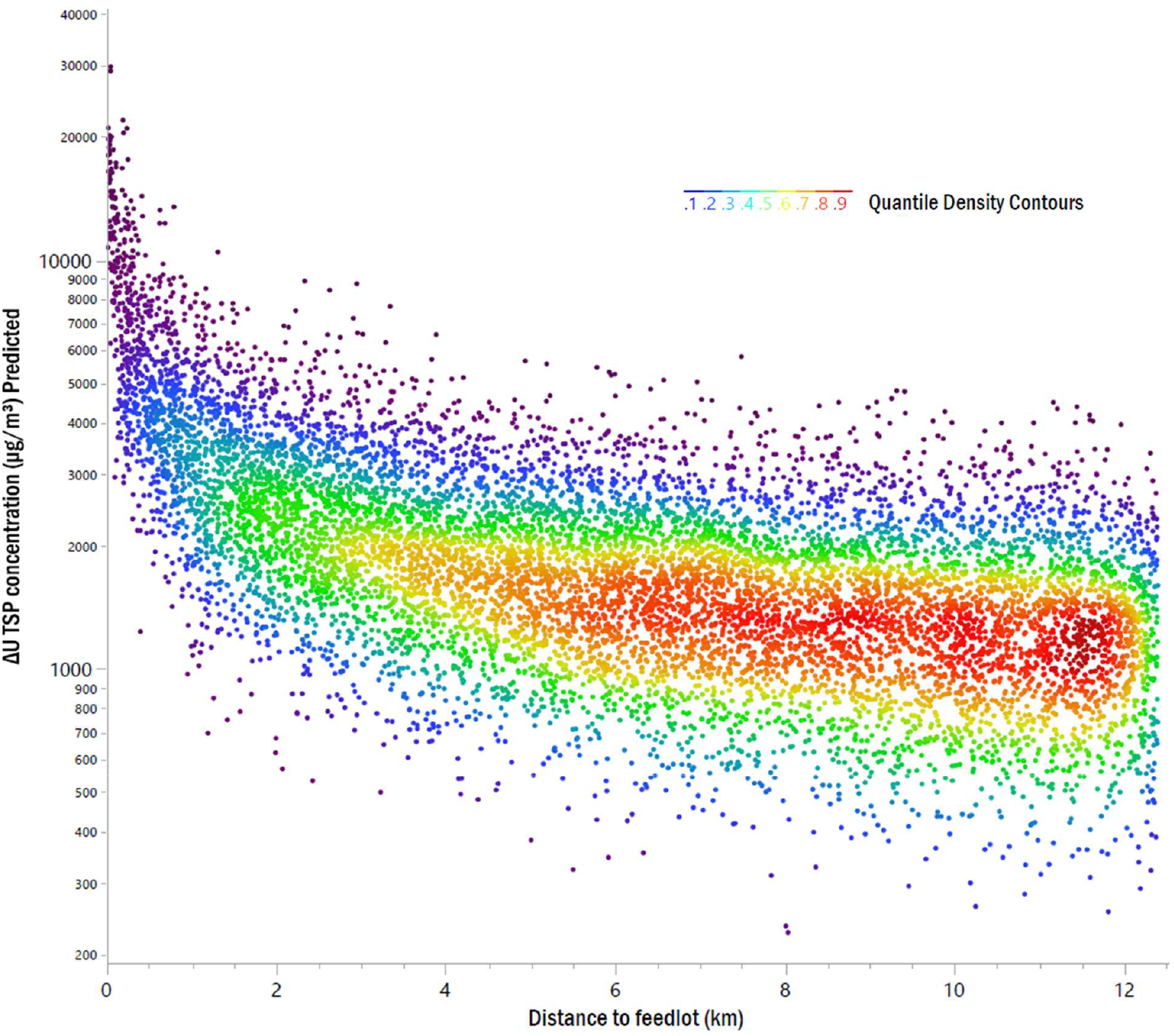
Monte Carlo simulation of feedlot-derived downwind ΔU TSP concentrations (µg/m^3^) downwind of three feedlots (F1-F3) in the Southern Great Plains, Texas, USA.

**Table 2:** Actual and Monte Carlo-simulated Weibull growth parameter estimates of downwind distance decay; simulated values derived from ambient air ΔU TSP concentrations (µg/m^3^) measured at three downwind (D1-D3) locations at three fee dlots in the Southern Great Plains, Texas, USA.

| Parameters | Box-cox[ΔU TSP (μg/m<sup>3</sup>); λ = -0.109] |  | ΔU TSP (μg/m<sup>3</sup>) | 95% CI |  |
| --- | --- | --- | --- | --- | --- |
|  | Actual Mean ±SEM | Simulated Mean ±SEM<sup>a</sup> | Simulated Mean ±SEM<sup>a</sup> | Lower | Upper |
| asymptote (a) | 43.9 ± 290.5 | 40.0 ± 14.1 | 46,423.2 ± 7,112.6 | 32,482.8 | 60,363.6 |
| inflection point (b) | 1.87x10<sup>-5</sup> ± 0.002 | 7.97x10<sup>-5</sup> ± 0.0004 | 0.004 ± 0.002 | 0.0007 | 0.008 |
| growth rate (c) | -0.095 ± 0.161 | -0.098 ± 0.009 | -0.45 ± 0.006 | -0.46 | -0.44 |

<sup>a</sup>Monte Carlo simulation; n = 10,000 SEM = standard error of the mean CI = confidence interval

Seasonal exposure to downwind PM is greater for residents living NNW-NNE of feedlot facilities in the Texas panhandle (**Supplemental Material SI7**). From 1 May – 31 August 2022, prevailing winds were observed from the SSW-SSE (40.8%) with 31.4% overall observations between 10.0-14.9 mph. Excluding wet days (n = 53; precipitation in the previous 72 hrs), 70 dry days were retained for expected seasonal agrochemical exposure frequency (EF).

### 3.2 Agrochemical occurrence

Of 11 agrochemical analytes included in the current study, four of six pyrethroids and all five MLs were detected in downwind TSP >LOQ (**Table 3**). Pyrethroids were detected at the highest overall concentrations >LOQ in 96.2% of downwind (grand mean = 3.4 ±0.5 ng/m^3^) and 27.8% of upwind samples (grand mean = 1.2 ±0.6 ng/m^3^), with fenvalerate detected at the highest mean downwind concentration (5.9 ± 0.8 ng/m^3^) followed by permethrin (1.1 ± 0.3 ng/m^3^). Bifenthrin and λ-cyhalothrin were detected infrequently >LOQ in 5.6 and 3.7% of all samples, respectively. Cypermethrin was not detected >LOQ in any samples, and esfenvalerate was detected only once in an upwind sample (F3; 2.8 ng/m^3^). At least one ML was detected in 98.1% of downwind (grand mean = 0.2 ±0.04 ng/m^3^) and 100.0% of upwind samples (grand mean = 0.1 ±0.04 ng/m^3^). Among MLs, eprinomectin was detected at the highest mean downwind concentrations >LOQ (0.4 ±0.2 ng/m^3^) followed by ivermectin (mean = 0.3 ±0.1 ng/m^3^). Median ± median absolute deviation of pyrethroid and ML concentrations (ng/m^3^) by upwind and downwind distance classes (D1-D3) can be found in **Supplemental Material SI8a-b**.

**Table 3:** Pyrethroid and macrocyclic lactone (ML) occurrence in total suspended particulates (TSP) upwind and downwind (D1-D3) of three feed lots in the Southern Great Plains, Texas, USA.

| Compound | Class | Detection (%)<sup>a</sup> | LOQ<sup>c</sup> | Recovery<br>(mean % ± SEM) | Mean<sup>ab</sup> | Median<sup>ab</sup> | Min.<sup>abc</sup> | Max.<sup>ab</sup> | SEM<sup>ab</sup> |
| --- | --- | --- | --- | --- | --- | --- | --- | --- | --- |
| bifenthrin<sup>d</sup> | Pyrethroid | 0 / 5.6 | 0.6 | 45.2 ± 4.5 | 0.5 | 0.5 | ND | 0.82 | 0.2 |
| λ-cyhalothrin<sup>d</sup> | Pyrethroid | 0 / 3.7 | 1.9 | 71.3 ± 5.4 | 0.9 | 0.9 | ND | 5.3 | 0.3 |
| cypermethrin<sup>d</sup> | Pyrethroid | 0 / 0 | 4.1 | 83.6 ± 2.8 | ND | - | - | - | - |
| esfenvalerate<sup>df</sup> | Pyrethroid | 5.6 / 0 | 2.8 | 68.4 ± 1.7 | 2.8 / 0 | - | - | - | - |
| fenvalerate<sup>d</sup> | Pyrethroid | 11.1 / 74.1 | 3.8 | 78.1 ± 10.1 | 0.2 / 5.9 | 0.2 / 4.3 | ND | 17.0 | 0.1 / 0.8 |
| permethrin<sup>d</sup> | Pyrethroid | 16.7 / 66.7 | 3.2 | 76.3 ± 8.5 | 1.3 / 1.1 | 0.5 / 0.7 | ND | 3.6 | 0.9 / 0.3 |
| abamectin<sup>e</sup> | ML | 50.0 / 48.1 | 0.2 | 74.5 ± 1.5 | 0.2 / 0.1 | 0.04 / 0.02 | ND | 1.0 | 0.1 / 0.05 |
| doramectin<sup>e</sup> | ML | 77.8 / 57.4 | 0.2 | 60.5 ± 7.8 | 0.07 / 0.07 | 0.04 / 0.03 | ND | 3.0 | 0.03 / 0.02 |
| eprinomectin<sup>e</sup> | ML | 33.3 / 33.3 | 0.2 | 80.5 ± 12 | 0.3 / 0.4 | 0.1 / 0.1 | ND | 3.0 | 0.2 / 0.2 |
| ivermectin<sup>e</sup> | ML | 11.8 / 66.7 | 0.2 | 90.0 ± 14.1 | 0.01 / 0.3 | 0.01 / 0.08 | ND | 2.7 | 0.01 / 0.1 |
| moxidectin<sup>e</sup> | ML | 88.9 / 81.5 | 0.2 | 70.6 ± 10.1 | 0.03 / 0.08 | 0.02 / 0.04 | ND | 0.9 | 0.01 / 0.02 |

Downwind summary statistics include all downwind locations (D1-D3; <1 – 12.4 km); six sampling events per feedlot; 30 min. duration per sample; sampling season between Julian days 137-229; <sup>a</sup>Values reported as upwind/downwind or downwind only, when applicable; <sup>b</sup>values reported in ng/m<sup>3</sup>; <sup>c</sup> values reported in ng/mL; <sup>d</sup> insecticide; <sup>e</sup> anthelmintic; <sup>f</sup> *S,S*-fenvalerate isomer; LOQ = limit of quantitation; ML = macrocyclic lactone; ND = not detected; SEM = standard error of mean

### 3.3 Parametric and non-parametric multivariate analyses

Multivariate analyses were performed to establish the underlying relationship between environmental predictor variables and downwind ambient air concentrations. PLS regression and CCA are constrained ordination methods that maximize explanatory covariance and correlation, respectively, between multiple predictor and dependent variables by projecting linear recombinations of original variables in factor space. Recombination in both cases serve to retain as much of the original variation in the dataset, while removing multicollinearity that may confound results. PLS regression was used as an initial method to parse out two levels of meteorological effects (diurnal and intraseasonal), and determine associations between each gradient individually. Results of initial PLS regression of meteorological variables are detailed in **Supplemental Material SI9-13**, where associated scores along the Factor 1 and 2 axes preserved the greatest variation from the original dataset (Factor 1, X = 50.2%, Y = 34.0%; Factor 2, X = 49.8%, Y = 6.2%). PLS factor 1 and 2 scores, hereafter “intraseasonal weather variation” and “diurnal weather variation,” respectively, were retained for use in CCA.

Final predictor variables included in CCA were downwind feedlot distance (km), intraseasonal variation, and diurnal variation, and response variables included ΔU TSP and ΔU ambient air concentrations of fenvalerate, permethrin, ivermectin, and moxidectin. Downwind concentrations, intraseasonal variation, and diurnal variation were not significantly different across feedlots (whole model F = 0.53; p = 0.715). Downwind concentrations were significantly correlated (eigenvalue = 1.48, canonical correlation = 0.772) with feedlot proximity (**Figure 4**; univariate F = 10.9; p <0.0001; fenvalerate > TSP > ivermectin > moxidectin > permethrin). Alternatively, intraseasonal and diurnal weather variation were not significant predictors across sampling events (intraseasonal F = 0.52, p = 0.759; diurnal F = 0.041, p = 0.907). Friedman’s non-parametric tests (**Supplemental Material SI14**) established significant differences in abamectin, doramectin, and eprinomectin concentrations across all downwind distance classes (D1-D3) relative to upwind. Abamectin and eprinomectin concentrations were significantly different across all feedlots, whereas a significant difference was observed in doramectin concentrations only between F1 and F2. However, due to low overall concentrations of ivermectin and moxidectin, and infrequent detections of abamectin, doramectin, and eprinomectin, MLs were excluded from cumulative exposure assessment.

**Figure 4:**
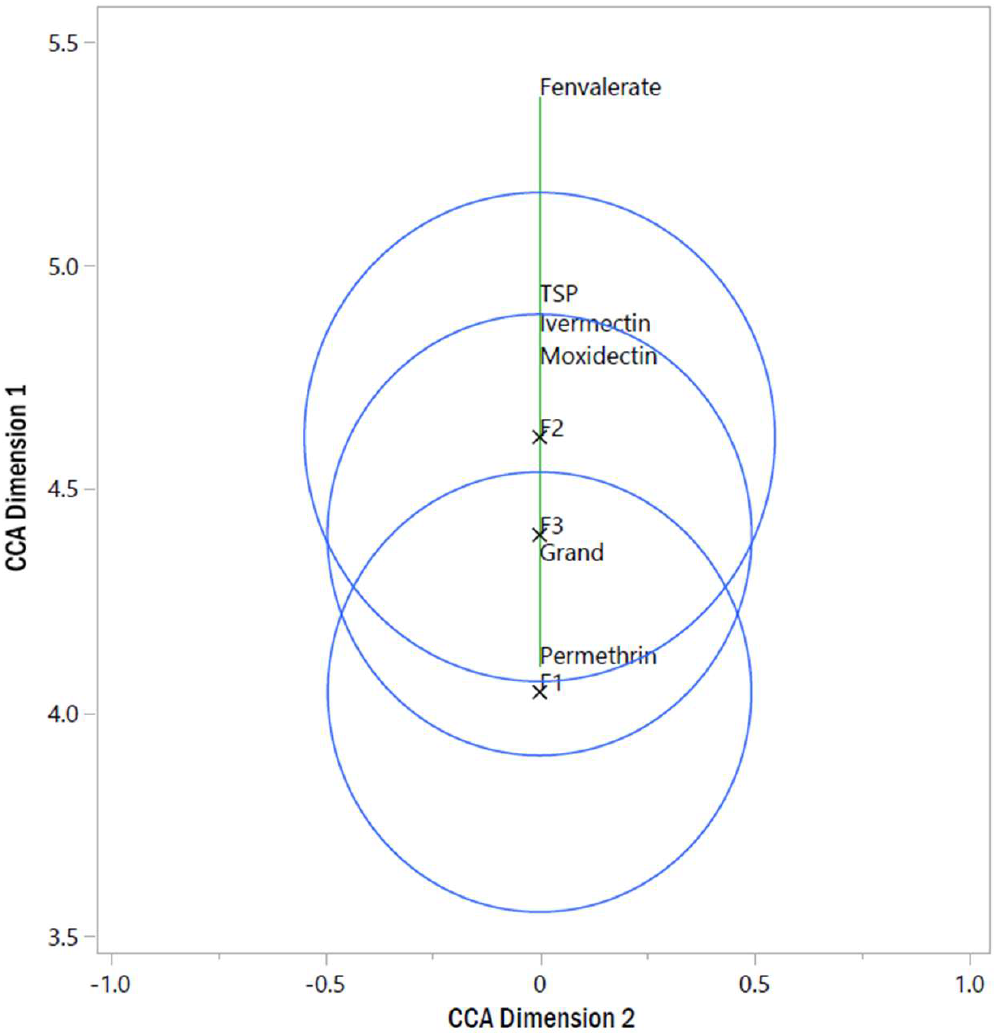
Canonical centroid plot (univariate: feedlot proximity) of ambient air concentrations downwind of three feedlots (F1-F3) in the Southern Great Plains, Texas, USA; canonical centroid values of individual consistuents in downwind ambient air are highlighted by overlapping biplot rays (green) across CCA dimension 1, indicating relative correlation magnitude with feedlot proximity; overlapping site centroids (blue) indicate no significant difference across feedlots; Grand = grand centroid value; TSP = total suspended particulates

**Figure 5:**
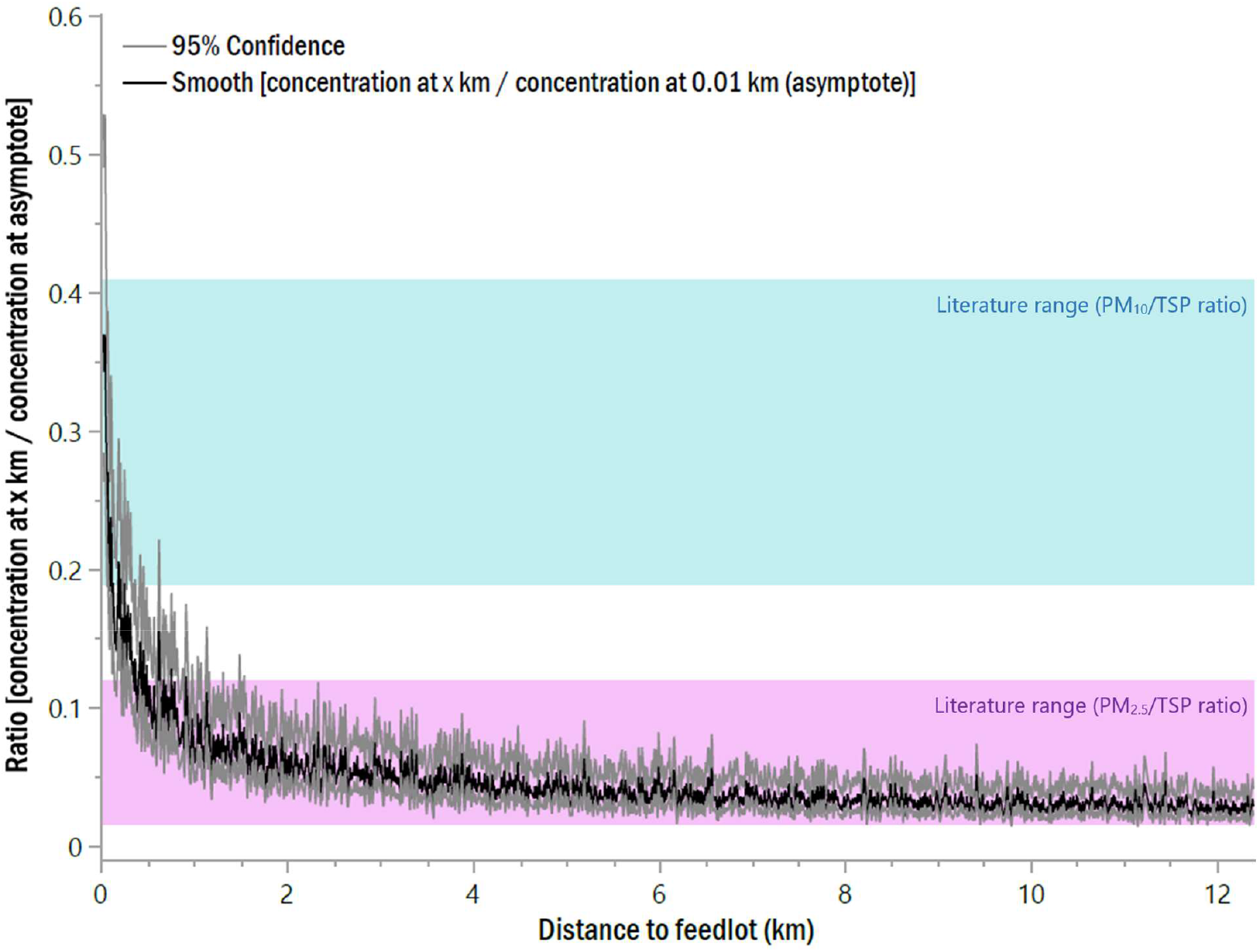
Moving 10-point average ratio of ΔU TSP ambient air concentrations *x* km downwind of feedlot divided by ΔU TSP ambient air concentrations at asymptote of Weibull decay curve (0.01 km); blue and purple shading highlights mean PM10/TSP and PM2.5/TSP ratios, respectively, reported within feedlot boundaries (Blackwell et al. 2015; Bonifacio et al. 2015) and downwind-adjacent to feedlots (Guo et al. 2011; Sweeten et al. 1998).

### 3.4 Cumulative human health assessment

A cumulative residential human health assessment was performed for pyrethroids, including feedlot-derived inhalation exposure estimates for fenvalerate and permethrin, due to frequency of occurrence and ambient air concentrations in the current study. Feedlot-derived ML inhalation exposure estimates (99.9^th^ percentiles) are also provided for reference in **Supplemental Material SI15**. Individual and aggregate MOEs and cumulative RIs for pyrethroids are reported in **Table 4**. Post-application edge-of-field exposure to aerially-sprayed bifenthrin (assumed wind vector = 10 mph) was selected as a protective residential exposure scenario due to frequent use as a foliar treatment for cotton leafhoppers (Vyavhare & Kerns, 2022), the most prevalent and expensive foliar cotton pest in the Texas Panhandle in 2021 (100% cotton acreage infested; 13% of Texas cotton acreage reported as insecticide-treated; Cook & Threet, 2021). The pyrethroid index POD (11 mg/kg/day) was derived from deltamethrin and adjusted based on compound-specific relative potency factors (RpF; U.S. EPA, 2011). All remaining exposure estimates were based directly on U.S. EPA 2011 residential and dietary exposure scenarios considered protective for cumulative exposure, except for residential handler and post-application exposure to backpack-sprayed turf treatment. Fluvinate, originally included in 2011, has since been discontinued, so the next highest turf exposure scenario was selected. Adult lifetime-adjusted average daily dose (mg/kg/day) and incremental lifetime aggregate cancer risk were also calculated for permethrin (**Table 5;** Q_i_* = 9.567E^-3^), recently designated as a likely human carcinogen (U.S. EPA, 2017a).

**Table 4:**
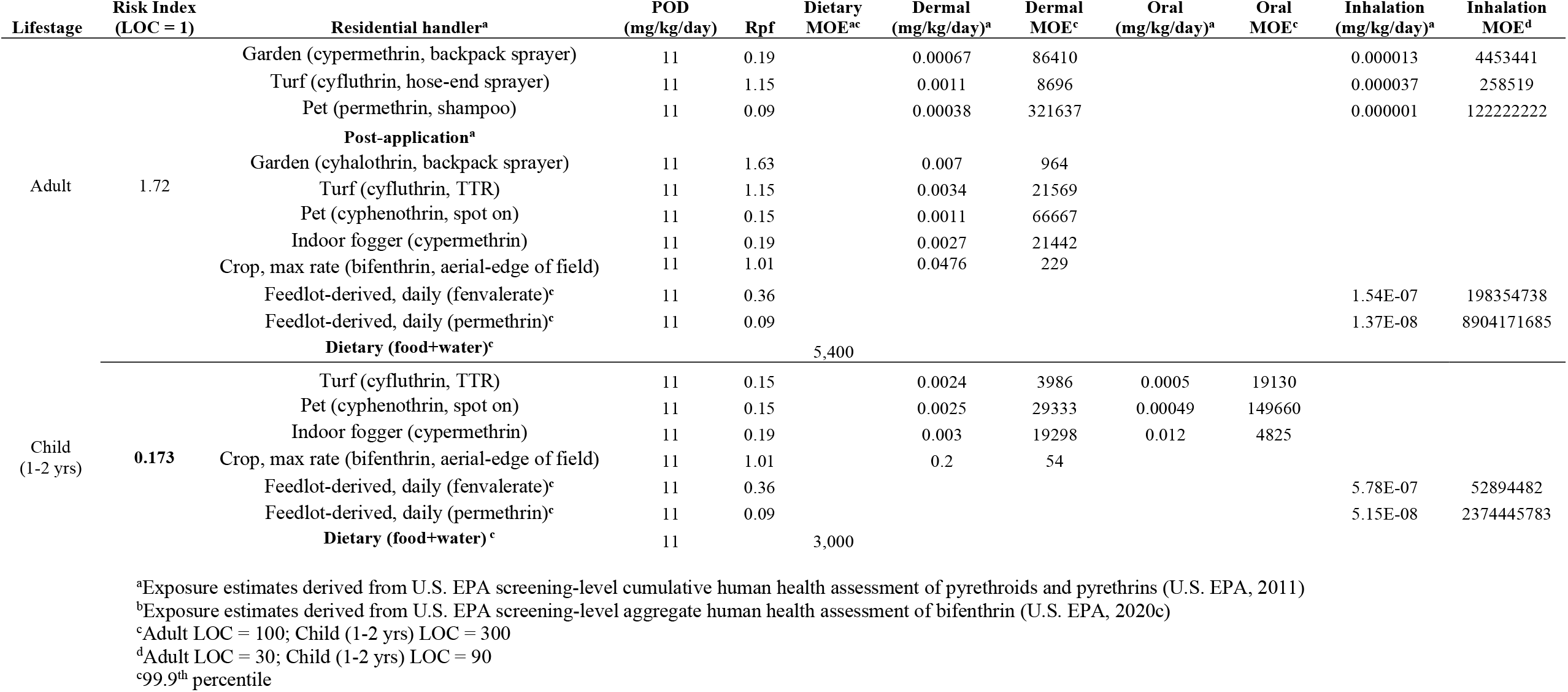
Cumulative non-carcinogenic human health assessment of pyrethroids in agricultural communities downwind of feedlots in the Southern Great Plains, Texas, USA.

**Table 5:** Adult residential feedlot-derived and aggregate lifetime cancer risk of permethrin in agricultural communities downwind of feedlots in the Southern Great Plains, Texas, USA.

| LADD (mg/kg/day;<br>feedlot-derived) | Cancer potency factor<br>($Q_i^*$)<sup>a</sup> | ILCR (feedlot-derived) | Aggregate ILCR<sup>a</sup> | |
| --- | --- | --- | --- | --- |
|  |  |  | Lower | Upper |
| 6.86E-10 | 9.57E-3 | 6.56E-12 | 3.83E-06 | 5.10E-05 |

<sup>a</sup>Aggregate values derived from ranges reported in U.S. EPA permethrin aggregate human health assessment (U.S. EPA, 2017a)
ILCR = incremental lifetime cancer risk

## 4. Discussion

### 4.1 Feedlot-derived PM

Particulate matter of feedlot origin has gained research traction in the last two decades, but considerable uncertainties remain regarding the full spatial extent and magnitude of atmospheric transport. Previous literature has indicated that TSP within and downwind-adjacent to feedlot pens is largely comprised of PM size fractions >10 µm (PM_2.5_/TSP = 0.017 - 0.12, PM_10_/TSP = 0.19 – 0.41; Blackwell, Wooten, et al., 2015; Bonifacio et al., 2015; J. M. Sweeten et al., 1998). As such, most pen floor material subject to initial suspension is not capable of sustained atmospheric transport. However, due to considerable suspended mass overall, elevated levels of inhalable PM size fractions are frequently observed. As highlighted in **Figure 4**, ratios of TSP concentrations ≥1.6 – 12.4 km vs. TSP concentrations at the asymptote (0.01 km; mean ratio = 0.039; 95% CI: 0.004 - 0.147) are in good agreement with feedlot PM_2.5_/TSP ratios reported in previous literature. Presently, downwind distance decay modeling of feedlot TSP and previous literature would suggest that non-inhalable and coarse particulate size fractions are primarily deposited ≤1.6 km downwind. Alternatively, this also suggests that downwind TSP concentrations ≥1.6 - 12.4 km (1,791 ± 9.9; 95% CI: 1,773 – 1,811 µg/m^3^) are predominately composed of feedlot-derived PM_2.5_ at levels that would significantly impact downwind residents during diurnal peak periods. Current estimates are also supported by Hiranuma et al. (2011), who found that ground-level (2-m height) PM_10_ concentrations measured 3.5 km downwind of a feedlot were reduced to approximately 8.5% of concentrations measured at the edge of cattle pens, whereas PM_2.5_ concentrations did not significantly decrease across the same distance. This size-dependent pattern of PM transport and deposition has been observed across various applications and has been used extensively to characterize PM transport potential from roadways (Karner et al., 2010). Several indirect methods have been recommended to approximate PM size distribution when direct methods are cost-prohibitive or otherwise not feasible, including use of emissions factors for animal waste-producing facilities based on total pen area (g/m^2^/d) and/or stocking density (g/m^2^/d-1000 head; U.S. EPA, 2022a), estimation of transportable fractions from livestock operations on a county-wide basis (Pouliot et al., 2011), and multipliers to approximate PM_2.5_ from PM_10_ concentrations (Pace, 2005). These estimation methods are particularly applicable to research in low-income, rural communities with disparate air quality monitoring infrastructure, as is the case in the current study.

While ambient air sampling in the current study was limited to TSP concentrations over a 3-hr time-from-sunset period given the concurrent objectives of quantifying agrochemical constituents and targeting peak diurnal emissions, no literature derived PM/TSP size ratio at any measured downwind distance would produce mean ambient air PM concentrations below National Ambient Air Quality Standards (NAAQS; U.S. EPA, 2023). Prior to 1987, primary TSP NAAQS were 260 µg/m^3^ (24-hr) and 75 µg/m^3^ (annual). From 1987 to present, primary (health-based) and secondary (public welfare) NAAQS have instead been based on inhalable PM size fractions, with primary standards set in 2012 as 35 µg/m^3^ (24-hr; one allowable exceedance of 98^th^ percentile) and 12.0 µg/m^3^ (annual; arithmetic mean) for PM_2.5_ and primary and secondary standards of 150 µg/m^3^ (24-hr) for PM_10_. In January 2023, new U.S. EPA guidelines were proposed (U.S. EPA, 2023), where primary PM_2.5_ standards would be reduced to a range between 9.0 - 10 µg/m^3^. Consequently, downwind residents who remain outdoors ≥30 mins. during diurnal peak periods are likely exposed to PM concentrations exceeding safe levels for human exposure.

### 4.2 Agrochemical occurrence

Trends of agrochemical occurrence in downwind PM are comparable to previous feedlot-adjacent sampling campaigns in the SGP region (Peterson et al., 2020). While usage of individual agrochemicals for beef cattle pest management will vary across feedlots, pyrethroids and MLs have been consistently detected in downwind PM. Upwind sampling was included in the current study to demonstrate comparative PM emissions and normalize downwind concentrations for analysis, a methodological approach commonly employed in feedlot sampling campaigns. However, trace agrochemicals detected upwind >LOQ in the current study should not be misinterpreted as a measure of regional background concentrations. Because co-located upwind samplers were deployed near cattle pens (<0.5 km), resuspension of previously deposited feedlot-derived PM must be factored into observations on the date of collection. While all pyrethroids of interest are used concurrently on croplands, abamectin is the only ML registered as an active ingredient for use in crop protection (U.S. EPA, 2017b; Vyavhare & Kerns, 2022), indicating that upwind detections of MLs are likely of feedlot origin. Thus, feedlot-derived agrochemical deposition also facilitates additional routes for daily human exposure, as accumulation of residues on outdoor residential areas increase the likelihood and magnitude of dermal, oral, and inhalation exposure. Because these additive exposure scenarios cannot be adequately characterized with the present study design, they have been excluded from subsequent exposure estimates, and quantification is recommended as a next step for future research.

### 4.3 Cumulative pyrethroid exposure in downwind agricultural communities

Cumulative RIs for pyrethroids indicate elevated health risk for children 1-2 yrs. of age (LOC = 1; RI = 0.173) living near agricultural operations in the SGP. While U.S. EPA cumulative exposure and risk estimates are intended to be extremely conservative in lower-tiered assessments, they are only as conservative as the exposure scenarios considered. Under U.S. EPA’s tiered-assessment framework, single compound aggregate RIs above LOCs under the most protective scenarios will initiate a higher-tiered, refined assessment. However, because residential assessments do not include post-application exposure from surrounding agricultural operations, nor synergistic toxicity in any exposure scenario, risk estimates assumed to be protective are inadequate for their intended purpose. Synergists such as piperonyl butoxide (PBO; Group C carcinogen; U.S. EPA, 2022b) are standard in pyrethroid product formulations (Gajendiran & Abraham, 2018), significantly increasing toxicity of individual active ingredients. Further, post-application inhalation exposure of pyrethroids have been assumed to be negligible beyond re-entry periods citing low volatilization and initial aerosol deposition (U.S. EPA, 2011). As demonstrated in the current study, these assumptions are incorrect when agrochemical residues are subject to post-application mechanical re-suspension from cattle trampling and/or farming equipment. Inhalation exposure and aerial dispersion are subsequently driven by wind transport and aerodynamic properties of particulates to which compounds are sorbed. Even as the present assessment elucidates unacceptable health risks in children 1-2 yrs. of age, it must be intimated that RI’s likely still underestimate worst-case scenario additive and synergistic pyrethroid risk to agricultural communities downwind of feedlot operations.

### 4.4 Incremental lifetime cancer risk of pyrethroids

Lifetime cumulative cancer risk from possible human carcinogens (bifenthrin and cypermethrin; U.S. EPA, 2022b), and the more recently classified likely carcinogenicity of permethrin are also not included in U.S. EPA’s most recent 2011 cumulative pyrethroid assessment. A Tier II epidemiology report was initiated in 2011 due to “moderately large” incidences of acute permethrin poisoning. In 2019, U.S. EPA summarily rejected any carcinogenic and non-carcinogenic health outcomes identified in epidemiologic studies (n = 62; U.S. EPA, 2019a) as lacking sufficient evidence to conclude a clear association or causal relationship with chronic, low-level pyrethroid exposure. Specifically, two studies investigated an association with pyrethroid exposure and incidence of acute lymphoid leukemia (ALL) and acute myeloid leukemia (AML) in Brazilian children (Ferreira et al., 2013; U.S. EPA moderate quality rank) and children residing in Shanghai, China (Ding et al., 2012; U.S. EPA low quality rank). Further epidemiological evidence of pyrethroid association with blood cancer precursors (MGUS; Hofmann et al., 2021) have since been reported. In concert with more recent prenatal *in vivo* studies (Priyanka et al., 2022; Song et al., 2022), conflicting conclusions on the weight of epidemiological evidence for health outcomes in children (Andersen et al., 2022), and prevalent incidences of acute permethrin exposure, the supporting literature for carcinogenic, reproductive, and neurodevelopmental pyrethroid risk to children deserves significantly greater regulatory consideration.

### 4.5 Feedlot-derived ML exposure

Macrocyclic lactones are not expected to pose an elevated human health based on minimal overlapping cropland use, but the indirect implication of ML resistance in soil-transmitted helminths also deserves careful consideration. Though outside of the present scope, atmospheric deposition of MLs may represent an environmental health risk to non-target invertebrates, particularly in soil microbial communities and dung beetles (Sands and Noll, 2022; de Souza and Guimarães 2022). Lastly, agrochemical occurrence in PM is not limited to pesticides, and the potential for chronic exposure to complex mixtures of endocrine-disrupting compounds, including veterinary pharmaceuticals and bioaerosols downwind of feedlot facilities is an additional public and environmental health concern that must be addressed and regulated holistically.

### 4.6 Future directions

U.S. EPA’s most recent risk evaluation of beef cattle feedlot facilities (U.S. EPA, 2004) does not incorporate foundational advancements in environmental impact research over nearly two decades. Federal exemptions have since been applied to all animal waste-producing facilities, with the most recent legislation in 2019 eliminating all hazardous air release reporting for beef cattle feedlots of any size (U.S. EPA, 2023a). The current regulatory landscape surrounding feedlot facilities in the U.S. has shifted the onus of proof to impacted populations, often disabling unincorporated rural communities from mounting viable evidence of environmental and public health impact. Agrochemical occurrence and distribution in downwind PM also underpins the considerable knowledge gap regarding human exposure near feedlot facilities. Feedlot-derived risk estimates of individual pyrethroid and ML concentrations in ambient air are intended to lay the groundwork for future research to expand across other relevant compound classes, use patterns, exposure routes, and seasons. Thus, the current assessment only includes inhalation exposure from a single operation isolated from neighboring feedlots and upwind resuspension of previous feedlot-derived residues, and risk estimates should be interpreted or reproduced as such. Actual year-round chronic exposure to feedlot-derived PM and agrochemicals, and associated human health risk of complex PM constituents, is expected to be significantly greater in areas where feedlot operations are heavily concentrated near municipalities. As agrochemicals registered for concurrent use in crop production already exert a chemical burden in agroecosystems, all relevant agricultural inputs must be considered to address cumulative daily intake. Residential post-application exposure estimates of bifenthrin near cropland, while referenced regarding cotton pest usage, are derived from the same aerial max application rate (0.30 lb a.i./ac) for all crops treated with bifenthrin, except citrus and tobacco. Considering variable acreage, risk characterizations can be reasonably integrated for residential exposure across dominant bifenthrin-treated crops relevant to multiple agroecosystems. However, daily intake of pyrethroids cannot be realistically predicted without comprehensive regulatory monitoring of fugitive feedlot emissions.

## 4. Conclusions

In this study, authors characterized agrochemical and PM transport downwind of feedlot facilities, significantly expanding the known distribution of feedlot-derived agrochemicals. Feedlot-derived PM concentrations in downwind ambient air reported in the current study likely exceed NAAQS thresholds of safe exposure to downwind residential communities, irrespective of agrochemical occurrence. Moreover, estimates of cumulative pyrethroid intake in downwind agroecosystems are well-above LOCs for children 1-2 yrs. of age. Due to the complex constituents of feedlot-derived PM, and extensive residential and commercial use of pyrethroid, potential non-carcinogenic and carcinogenic health risks to residents warrant more comprehensive, transparent, and independent research. Aerial dispersion of PM-bound agrochemicals from feedlots is a major unrecognized transport pathway in environmental and human health assessments, and lack of monitoring and reporting of hazardous substance air releases at feedlots poses considerable challenges for risk characterization to sensitive receptors.

## Acknowledgement

This work was supported by the Texas Tech University Graduate School, the Texas Academy of Science Graduate Research Award, and the Sandy Land Underground Water Conservation Research Scholarship.

## Data Statement

Location-censored raw data tables are available by request to the corresponding author.

## Acronyms

AICc: Corrected Akaike’s information criterion for small sample sizes
FQPA: Food Quality Protection Act
FQPA SF: FQPA safety factor
GABA: gamma (γ)-aminobutryic acid
IRIS: Integrated Risk Information System, U.S. EPA
LOC: level of concern
LOD: limit of detection LOQ – limit of quantitation
MGUS: monoclonal gammopathy of undetermined significance
MADL: maximum allowable dose level; Proposition 65, California Office of Environmental Health Hazard Assessment
Min.: minimum
Max.: maximum
MOA: mechanism of action
MOE: margin of exposure
MSE: mean square error
NAAQS: National Ambient Air Quality Standards
NOAEL: No-observed-adverse-effect level
PK: pharmacokinetic
PM: particulate matter
PM_2.5_: particulate matter of aerodynamic equivalent diameter ≤2.5 μm
PM_10_: particulate matter of aerodynamic equivalent diameter ≤10 μm
POD: point-of-departure
RpF: relative potency factor
RMSE: root mean square error
SEM: standard error of the mean
SD: standard deviation
SSE: sum of squares error
U.S. EPA: United States Environmental Protection Agency
OPP: U.S. EPA Office of Pesticide Programs
SOP: standard operating procedure
USDA: United States Department of Agriculture
NASS: USDA National Agricultural Statistics Service
TSP: total suspended particulates
ΔU: upwind-normalized
UF_A_: uncertainty factor, interspecies extrapolation
UF_H_: uncertainty factor, interindividual variability (human)

