## Supplemental Material for "Atmospheric transport of particulate matter and particulate-bound agrochemicals from beef cattle feedlots: human health implications for downwind agricultural communities"

**SI1:** Site characteristics of three feedlots (F1-F3) sampled in the Southern Great Plains, Texas, USA

| Site characteristic | F1 | F2 | F3 |
|---|---|---|---|
| Animal unit capacity (head) | 75,000 | 82,500 | 80,000 |
| Pen area (ac) | 149 | 201 | 271 |

ac = acre

**SI2:** Mahalanobis' distance (n = 5) and ΔU TSP (μg/m³) residual quantile outliers (n = 2)

| Outliers (Feedlot/Sample ID) | Julian day | ΔU TSP (μg/m³) | Distance (UCL = 4.38[a]) |
|---|---|---|---|
| F1, D1 | 207 | 80,817.5 | 4.53 |
| F2, D1 | 142 | 20,819.3 | 4.97 |
| F3, D1 | 165 | 3,353.1 | 4.90 |
| F2, D2[b] | 165 | 209.1 | 3.46 |
| F2, D3 | 142 | 0.00 | 4.77 |
| F2, D3[b] | 149 | 3.11 | 4.19 |
| F3, D3 | 210 | 0.00 | 4.74 |

[a]UCL = upper control limit
[b]Residual quantile outlier (>4 standard deviations)

**SI3**: Descriptive statistics and feedlot proximity (km) of total suspended particulates (TSP) samples collected at one upwind and three downwind (D1-D3) locations at three feedlots (F1-F3) in the Southern Great Plains (SGP), Texas, USA.

| | | | **TSP (μg/m³)** | | | | | | **Distance to feedlot (km)** | | | | | |
|---|---|---|---|---|---|---|---|---|---|---|---|---|---|---|
| Site | Sample ID | n[a] | Mean | Median | SD | SEM | Min. | Max. | Mean | Median | SD | SEM | Min. | Max. |
| F1 | Upwind | 18 | 116.3 | 131.2 | 56.3 | 17.8 | 47.5 | 189.5 | 0.46 | 0.47 | 0.05 | 0.02 | 0.35 | 0.48 |
| | D1 | 18 | 36604.1 | 23433.1 | 28502.0 | 8593.7 | 6599.0 | 83321.6 | 0.02 | 0.02 | 0.004 | 0.002 | 0.02 | 0.03 |
| | D2 | 18 | 3017.7 | 2706.5 | 1580.5 | 476.5 | 1362.1 | 5621.4 | 3.22 | 3.30 | 0.20 | 0.08 | 2.84 | 3.46 |
| | D3 | 18 | 1572.2 | 2223.3 | 934.3 | 281.7 | 462.5 | 2527.2 | 10.11 | 10.20 | 1.21 | 0.49 | 8.09 | 11.86 |
| F2 | Upwind | 18 | 457.8 | 225.0 | 664.1 | 210.0 | 39.2 | 2376.8 | 0.06 | 0.05 | 0.02 | 0.01 | 0.03 | 0.11 |
| | D1 | 18 | 19944.7 | 19788.3 | 8986.3 | 2709.5 | 6029.8 | 34493.0 | 0.06 | 0.04 | 0.07 | 0.03 | 0.01 | 0.21 |
| | D2 | 18 | 2504.6 | 2130.9 | 1373.8 | 414.2 | 421.7 | 5246.5 | 3.53 | 3.49 | 1.15 | 0.47 | 1.97 | 5.17 |
| | D3 | 18 | 1270.1 | 1577.1 | 740.0 | 223.1 | 62.7 | 1892.3 | 10.44 | 10.80 | 1.72 | 0.70 | 7.28 | 12.43 |
| F3 | Upwind | 18 | 213.8 | 163.4 | 154.7 | 48.9 | 57.0 | 605.2 | 0.07 | 0.07 | 0.04 | 0.02 | 0.03 | 0.10 |
| | D1 | 18 | 4710.2 | 3560.2 | 3129.3 | 943.5 | 1937.2 | 11470.7 | 0.76 | 0.85 | 0.35 | 0.14 | 0.12 | 1.09 |
| | D2 | 18 | 3458.3 | 2989.0 | 2676.7 | 807.1 | 929.3 | 10467.5 | 4.31 | 4.10 | 1.22 | 0.50 | 3.13 | 5.88 |
| | D3 | 18 | 1730.9 | 1684.8 | 1280.3 | 386.0 | 291.2 | 3579.4 | 10.78 | 10.30 | 1.05 | 0.43 | 9.72 | 12.39 |

[a]Six sampling events per feedlot; 30 min. duration per sample; sampling season between Julian days 137-229
μg/m³ = microgram per cubic meter
km = kilometer
SD = standard deviation
SEM = standard error of the mean
Min. = minimum
Max. = maximum

**SI4:** Empirical ΔU TSP (μg/m³) distance decay model comparison of training/validation set

| Model[a] | ΔAICc | AICc weight (ωi) | SSE | MSE | RMSE | $R^2$ |
|---|---|---|---|---|---|---|
| Weibull Growth | - | 0.4713 | 111.0 | 3.70 | 1.92 | 0.588 |
| Biexponential 4p | 0.55 | 0.3579 | 103.7 | 3.58 | 1.89 | 0.615 |
| Biexponential 4p | 3.40 | 0.0862 | 103.2 | 3.68 | 1.92 | 0.617 |
| Exponential 3P | 3.51 | 0.0814 | 123.4 | 4.11 | 2.03 | 0.542 |
| Exponential 2P | 10.4 | 0.0027 | 164.3 | 5.30 | 2.30 | 0.390 |
| Logistic 3P | 13.0 | 0.0007 | 164.3 | 5.48 | 2.34 | 0.390 |

[a]Model comparison derived from training/validation set

**SI5a-b:** Weibull-distributed a) actual vs. predicted and b) residual vs. predicted box-cox-transformed ΔU TSP (ug/m³) plots of training/validation set

**a)**

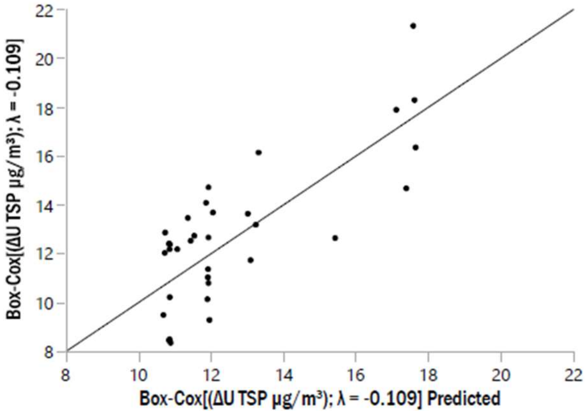


**b)**

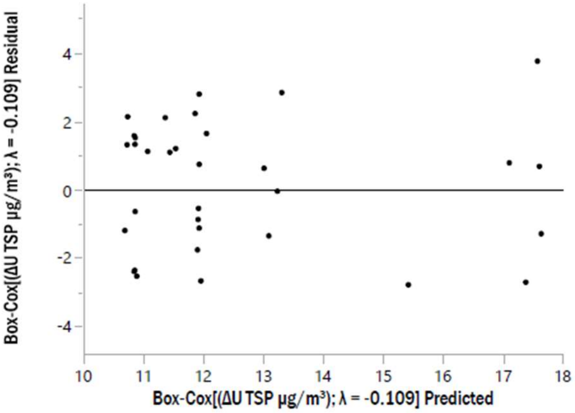


**SI6a-b:** Weibull-distributed ΔU TSP distance decay a) training and validation model curve fit parameters and b) training/validation, test, and overall linear fit of transformed actual vs. predicted ΔU TSP concentrations

**a)**

| **Weibull fit parameters** | Training | Validation | Training/validation |
|---|---|---|---|
| Sum of squares error | 62.7 | 42.9 | 111.0 |
| Mean square error | 4.48 | 3.30 | 3.70 |
| Root mean square error | 2.12 | 1.82 | 1.92 |
| $R^2$ | 0.60 | 0.62 | 0.59 |

**b)**

| **Linear fit parameters (actual v. predicted)** | Train/Validate | Test | Overall |
|---|---|---|---|
| $R^2$ | 0.59 | 0.95 | 0.71 |
| df | 32 | 13 | 45 |
| Sum of squares error | 55.3 | 3.81 | 63.6 |
| Mean square error | 1.79 | 0.32 | 1.41 |
| F-ratio | 44.5 | 235.9 | 108.6 |
| p-value | <0.0001* | <0.0001* | <0.0001* |

**SI7:** Collated wind rose (n = 4 stations) from 1 May – 31 August 2022 highlighting prevailing cardinal wind direction and wind speed (mph) at three feedlots (F1-F3) in the Southern Great Plains, Texas, USA

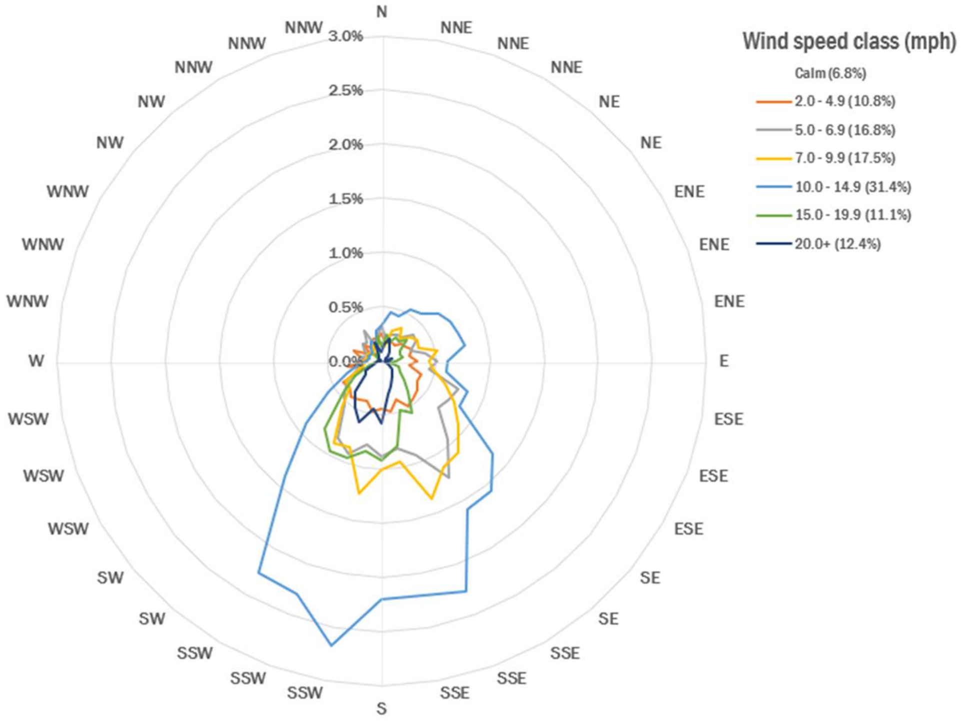

**SI8a-b:** Median ± median absolute deviation a) pyrethroid and b) macrocyclic lactone (ML) concentrations (ng/m$^3$) measured at one upwind and three downwind locations (D1-D3) at three feedlots (F1-F3; six evenings per feedlot) in the Southern Great Plains, Texas, USA; non-detect median for bifenthrin, esfenvalerate, and λ-cyhalothrin

**SI8a)**

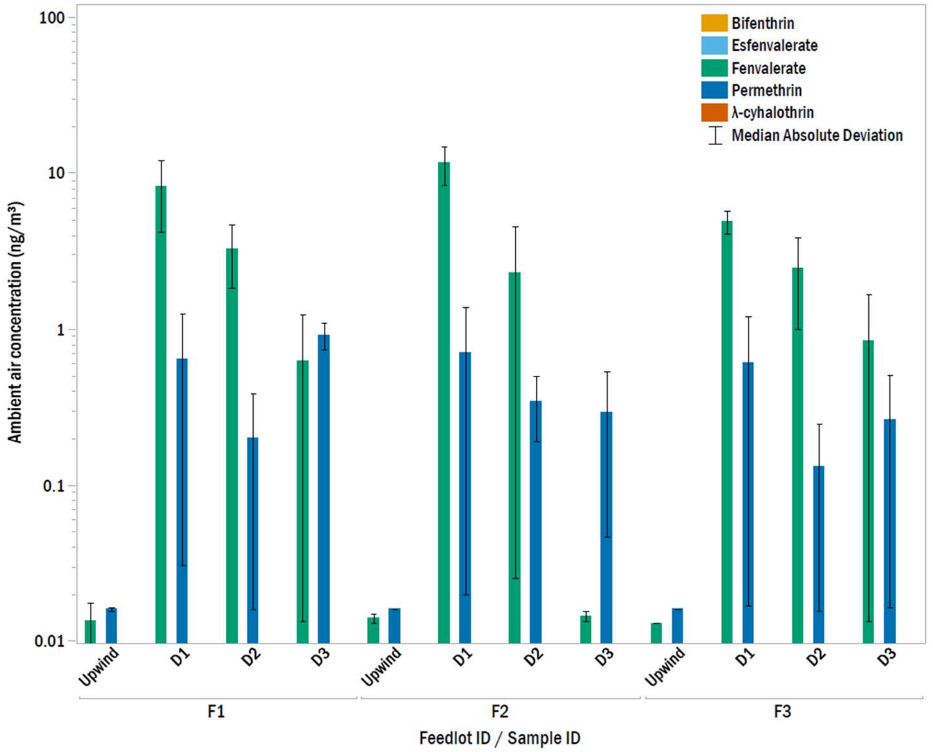


**SI8b)**

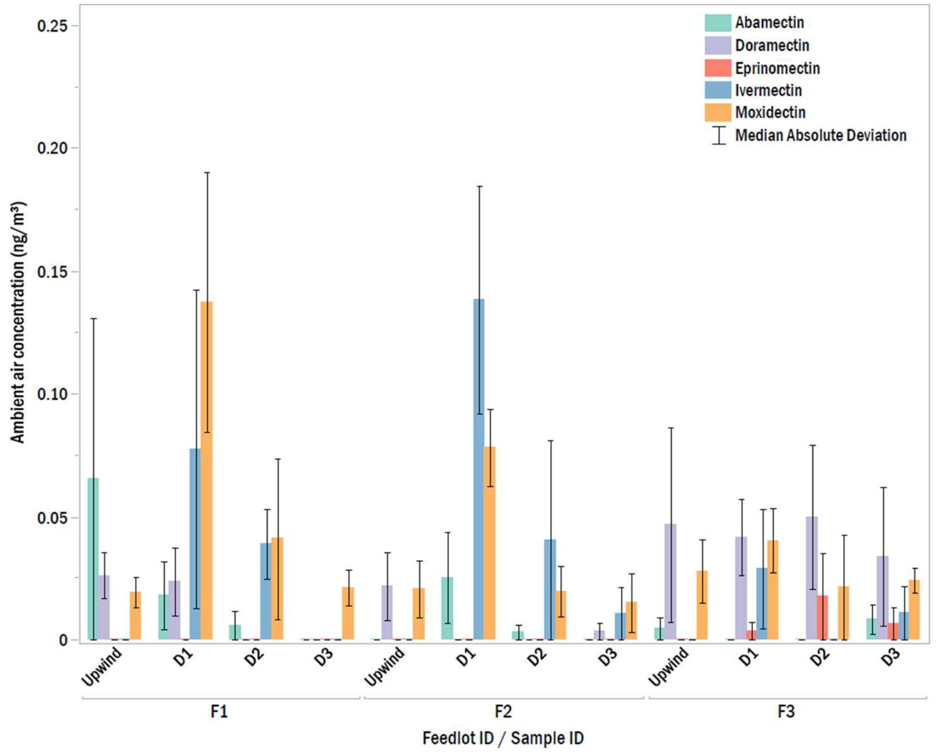

**SI9: Partial least squares (PLS)**

All measured weather variables (Y responses; **SI10**) constrained to Julian day and time from sunset (X effects), were included in PLS. Two factors (root mean PRESS = 0.90) explained a cumulative 100.0 and 40.2% of variation in X and Y, respectively (**SI11**), where intraseasonal variation was maximized on Factor 1 and diurnal weather variation was maximized on Factor 2. Variable importance was significant (**SI12;** VIP ≥0.8) for julian day (VIP = 1.41), but was not significant for time from sunset (VIP = 0.11) indicating relatively low variation across diurnal weather gradients. Individually, however, wind speed exhibited the greatest variation between downwind distances (33.7%) with little variation across sampling events (5.93%; **SI13a-c**). Regardless of low diurnal variation, both factors have been included in subsequent analysis. Low variations in predictors examined independently do not necessarily indicate that potentially relevant associations are not present when constrained to ambient air concentrations in CCA. Thus, both Factor 1 and 2 scores were retained to preserve maximum variation of recombined variables.

**SI10:** Descriptive statistics of weather variables recorded by West Texas Mesonet (WTM) at one upwind and three downwind (D1-D3) sampling locations at three feedlots (F1-F3)

| Weather variable | n[a] | Mean | Median | STD | SEM | Min. | Max. |
|---|---|---|---|---|---|---|---|
| Wind speed (mph) | 72 | 11.9 | 11.5 | 3.0 | 0.4 | 7 | 23.0 |
| Wind direction (deg) | 72 | 157.8 | 157.5 | 38.2 | 4.5 | 67.5 | 247.5 |
| Temperature (°C) | 72 | 33.4 | 34.2 | 4.3 | 0.5 | 17.1 | 39.2 |
| Relative humidity (%) | 72 | 22.1 | 22.1 | 8.6 | 1.0 | 6.6 | 42.0 |
| Barometric pressure (inHg) | 72 | 26.7 | 26.8 | 0.2 | 0.02 | 26.1 | 26.9 |

[a]Six events per feedlot; sampling season between Julian days 137-229

**SI111:** Partial least squares (PLS) regression of measured meteorological variables; PLS Factor 1 and 2 scores included in canonical correlation analysis (CCA)

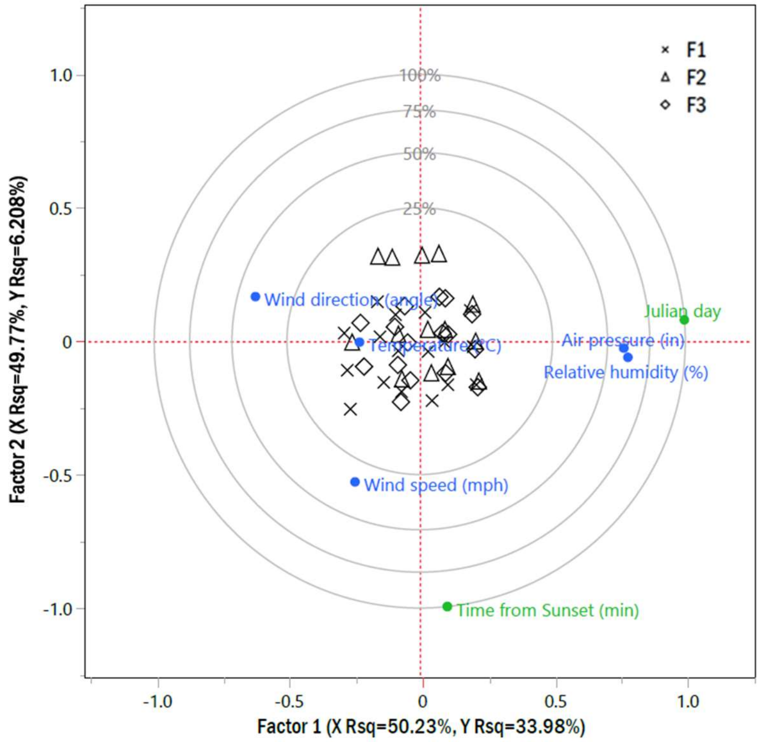

**SI12:** Partial least squares variable importance plot (VIP) of X effects (time from sunset and Julian day) to variation in Y responses (meteorological variables)

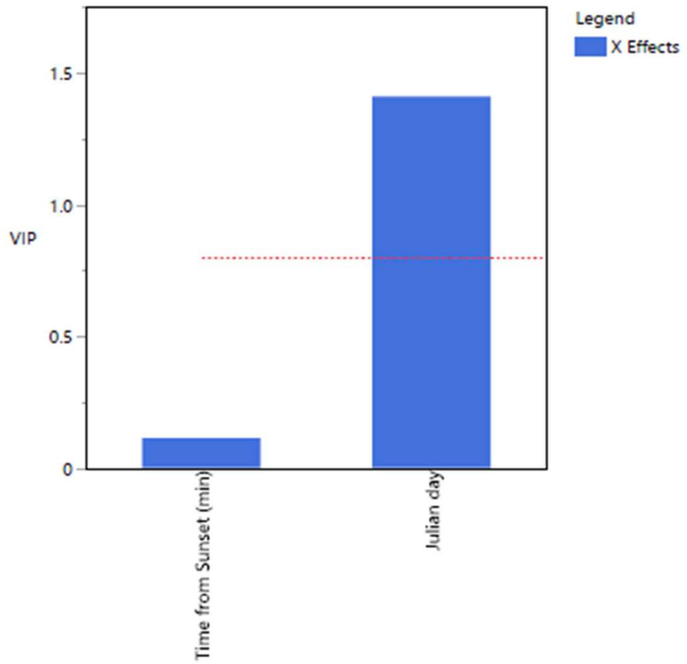


**SI13a-c:** Partial least squares percent variation explained for a) X effects (time from sunset and Julian day) and b/c) Y responses (relative humidity, air temperature and pressure, and wind speed and direction)

**SI13a)**

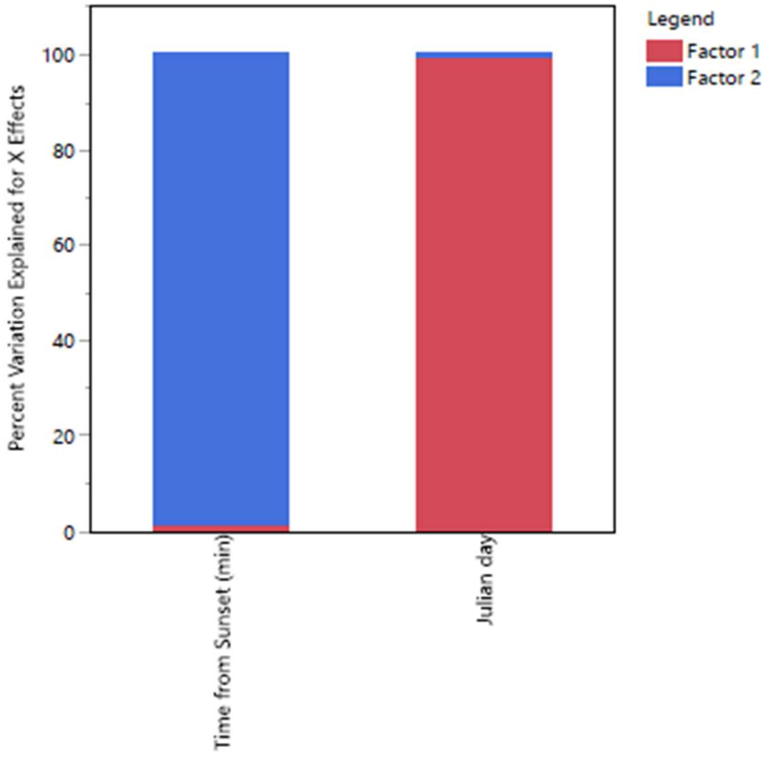

**SI13b)**

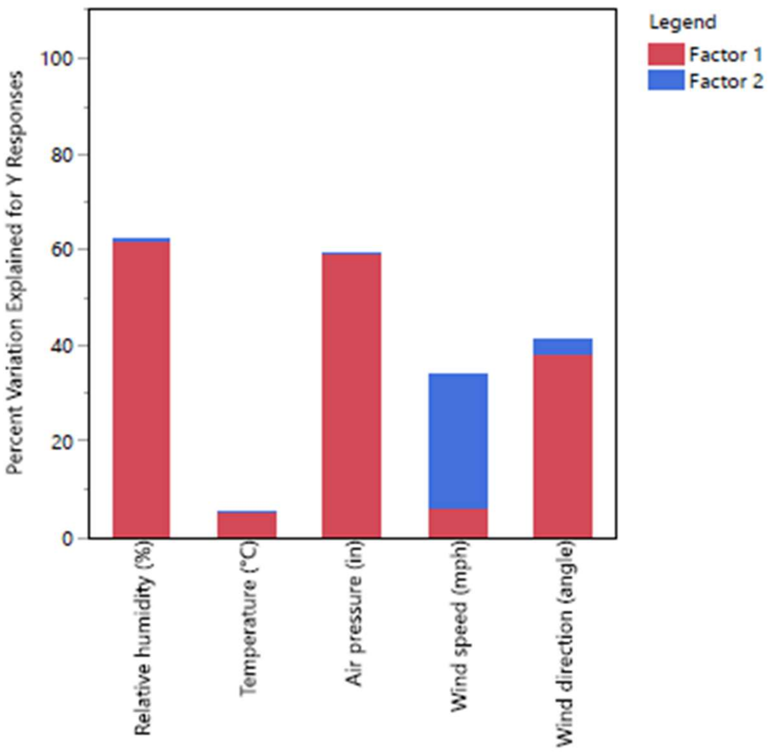


**SI13c**) Percent variation in Y responses explained by latent factors

| Y response | Factor 1 (%) | Factor 2 (%) |
|---|---|---|
| Relative humidity (%) | 61.5 | 61.9 |
| Temperature (°C) | 5.13 | 5.13 |
| Air pressure (in Hg) | 59.1 | 59.1 |
| Wind speed (mph) | 5.93 | 33.7 |
| Wind direction (angle) | 38.3 | 41.1 |

**SI14:** Friedman's non-parametric tests for abamectin, doramectin, and eprinomectin ambient air concentrations ($ng/m^3$) collected at one upwind and three downwind (D1-D3) locations at three feedlots (F1-F3) in the Southern Great Plains (SGP) region, Texas, USA

| Sample 1 v Sample 2; Analyte | Test Statistic | Std. Error | Std. Test Statistic | Sig. | Adj. Sig.[a] |
|---|---|---|---|---|---|
| Upwind v ΔU D1; abamectin ($ng/m^3$) | -3.056 | 0.624 | -4.900 | 0.000 | <0.001 |
| Upwind v ΔU D2; abamectin ($ng/m^3$) | -3.444 | 0.624 | -5.523 | 0.000 | <0.001 |
| Upwind v ΔU D3; abamectin ($ng/m^3$) | -2.500 | 0.624 | -4.009 | 0.000 | 0.001 |
| Upwind v ΔU D1; doramectin ($ng/m^3$) | -3.056 | 0.624 | -4.900 | 0.000 | <0.001 |
| Upwind v ΔU D2; doramectin ($ng/m^3$) | -3.444 | 0.624 | -5.523 | 0.000 | <0.001 |
| Upwind v ΔU D3; doramectin ($ng/m^3$) | -2.500 | 0.624 | -4.009 | 0.000 | 0.001 |
| Upwind v ΔU D1; eprinomectin ($ng/m^3$) | -3.056 | 0.624 | -4.900 | 0.000 | <0.001 |
| Upwind v ΔU D2; eprinomectin ($ng/m^3$) | -3.444 | 0.624 | -5.523 | 0.000 | <0.001 |
| Upwind v ΔU D3; eprinomectin ($ng/m^3$) | -2.500 | 0.624 | -4.009 | 0.000 | 0.001 |
| F1 v F2 - ΔU eprinomectin ($ng/m^3$) | 1.265 | 0.261 | 4.851 | 0.000 | <0.001 |
| F1 v F2 - ΔU abamectin ($ng/m^3$) | 2.633 | 0.261 | 10.094 | 0.000 | <0.001 |
| F1 v F2 - ΔU doramectin ($ng/m^3$) | 0.714 | 0.261 | 2.739 | 0.006 | 0.037 |
| F1 v F3 - ΔU eprinomectin ($ng/m^3$) | 1.125 | 0.264 | 4.269 | 0.000 | <0.001 |
| F1 v F3 - ΔU abamectin ($ng/m^3$) | 2.521 | 0.264 | 9.566 | 0.000 | <0.001 |
| F1 v F3 - ΔU doramectin ($ng/m^3$) | 0.604 | 0.264 | 2.293 | 0.022 | 0.131 |
| F2 v F3 - ΔU eprinomectin ($ng/m^3$) | 0.958 | 0.264 | 3.637 | 0.000 | 0.002 |
| F2 v F3 - ΔU abamectin ($ng/m^3$) | 2.438 | 0.264 | 9.250 | 0.000 | <0.001 |
| F2 v F3 - ΔU doramectin ($ng/m^3$) | 0.438 | 0.264 | 1.660 | 0.097 | 0.581 |

Each row tests the null hypothesis that the Sample 1 and Sample 2 distributions are the same; asymptotic significances (2-sided tests) are displayed; $\alpha = .05$; [a]Significance values have been adjusted by the Bonferroni correction for multiple tests; ΔU = upwind-normalized; SGP = Southern Great Plains

**SI15:** Downwind beef cattle feedlot-derived short-term macrocyclic lactone inhalation exposure estimates (99.9$^{th}$ percentile)

| Lifestage | Macrocyclic lactone | POD (mg/kg/day)[a] | Inhalation (mg/kg/day)[b] | Inhalation MOE[c] |
|---|---|---|---|---|
| Adult | abamectin | 0.25 | 7.66E-9 | 3.26E+07 |
| | doramectin | 0.25 | 1.96E-9 | 1.28E+08 |
| | eprinomectin | 0.25 | 2.28E-8 | 1.10E+07 |
| | ivermectin | 0.25 | 1.65E-8 | 1.52E+07 |
| | moxidectin | 0.25 | 1.97E-9 | 1.27E+08 |
| Child (1-2 yrs) | abamectin | 0.25 | 2.87E-8 | 8.71E+06 |
| | doramectin | 0.25 | 7.34E-9 | 3.41E+07 |
| | eprinomectin | 0.25 | 9.56E-8 | 2.62E+06 |
| | ivermectin | 0.25 | 6.18E-8 | 4.05E+06 |
| | moxidectin | 0.25 | 7.37E-9 | 3.39E+07 |

[a]Point-of-departure (POD; mg/kg/day) for all macrocyclic lactones derived from abamectin no-observed-adverse-effect level (NOAEL) reported in avermectin cumulative screening-level human health assessment (U.S. EPA, 2017b)
[b]99.9$^{th}$ percentile; downwind 0 – 12.4 km
[c]MOE = margin of exposure; LOC = 30; Child (1-2 yrs) LOC = 90
